# 3-phenyllactic acid promotes healthspan through SKN-1/ATFS-1-mediated mitochondrial activation and stress resilience

**DOI:** 10.1101/2023.11.13.566789

**Authors:** Juewon Kim, Yunju Jo, Gyumin Lim, Yosep Ji, Jong-Hwa Roh, Wan-Gi Kim, Hyon-Seung Yi, Dong Wook Choi, Donghyun Cho, Dongryeol Ryu

## Abstract

The mechanisms underlying the impact of probiotic supplementation on health remain largely elusive. While previous studies have primarily focused on the discovery of novel bioactive bacteria and alterations in the microbiome environment to explain potential probiotic effects, our research delves into the role of living lactic acid bacteria (LAB) and their conditioned media, highlighting that only the former, not dead bacteria, enhance the healthspan of *Caenorhabditis elegans*. To elucidate the underlying mechanisms, we conducted transcriptomic profiling through RNA-seq analysis in *C. elegans* exposed to LAB or 3-phenyllactic acid (PLA), mimicking the presence of key candidate metabolites of LAB and evaluating healthspan. Our findings reveal that PLA treatment significantly extends the healthspan of *C. elegans* by promoting energy metabolism and stress resilience in a SKN-1/ATFS-1-dependent manner. Moreover, PLA-mediated longevity is associated with a novel age-related parameter, the Healthy Aging Index (HAI), introduced in this study, which comprises healthspan-related factors such as motility, oxygen consumption rate (OCR), and ATP levels. Extending the relevance of our work to humans, we observed an inverse correlation between blood PLA levels and physical performance in patients with sarcopenia, when compared to age-matched non-sarcopenic controls. Our investigation thus sheds light on the pivotal role of the metabolite PLA in probiotics-mediated enhancement of organismal healthspan, and also hints at its potential involvement in age-related sarcopenia. These findings warrant further investigation to delineate PLA’s role in mitigating age-related declines in healthspan and resilience to external stressors.

**Highlights:**

- Metabolites Produced by Lactic Acid Bacteria Function as Mediators of Longevity in *C. elegans*
- 3-Phenyllactic Acid (PLA) Augmented Lifespan Extension and Elevated the Healthy Aging Index (HAI)
- PLA Facilitated Healthy Aging by Inducing Mitochondrial Activation and Stress Resistance Dependent on SKN-1/ATFS-1
- Patients with Sarcopenia and Symptoms of Frailty Exhibited Reduced Blood PLA Levels.

## Introduction

Diet plays an indispensable role in determining an organism’s health status, and consequently, it significantly impacts longevity. Over the years, numerous studies have concentrated on dietary components, nutrients, and ingredients that possess the potential to retard the aging process in invertebrate model organisms and mammals, including humans ^1–3^. Among these dietary components, lactic acid bacteria (LAB) have emerged as noteworthy candidates. LAB, classified as microorganisms that produce lactic acid as an end product of carbohydrate fermentation, have been shown to extend the lifespan of aging model organisms and enhance various health-related phenotypes during the aging process ^4–7^. While much research has been dedicated to exploring the anti-aging properties of LAB, investigations into the specific metabolites or ingredients responsible for these effects remain limited. LAB are living microorganisms that contribute to the restoration of intestinal microbial balance, immune modulation, and the exertion of probiotic activities within host organisms. Consequently, it is reasonable to assume that certain metabolites derived from LAB have the potential to influence numerous physiological changes through dietary supplementation. As a result, we have formulated a hypothesis that the increased metabolites resulting from interventions with LAB might have an impact on age-related health factors. To substantiate this conjecture, we conducted experiments to validate the longevity effects of LAB under various conditions using the well-established longevity model organism, *Caenorhabditis elegans* (*C. elegans*), known for its highly conserved metabolism. Our investigations confirmed that both the living state of LAB and the conditioned media of LAB could exert anti-aging effects, with the most notable increase being in the metabolic intermediate 3-phenyllactic acid (PLA), which we have identified as the active metabolite responsible for mitigating age-associated declines and extending healthspan.

In this study, we propose that the anti-aging effects are primarily attributed to the living state of LAB, which enhances *C. elegans* healthspan by stimulating the production of LAB-derived PLA. While these initial findings underscore the significance of PLA, its specific roles and mechanistic pathways in the aging process largely remain uncharted. To establish whether the efficacy of LAB metabolites in influencing the aging process is a conserved trait across species, we investigated whether the alteration in lifespan induced by PLA could also be observed in the representative aging model, *C. elegans*. Our study sheds light on the pivotal role of PLA, a LAB-derived metabolite, in modulating *C. elegans* lifespan and healthspan. We propose that this modulation is mediated through the activation of SKN-1/NRF2 and the activating transcription factor associated with stress-1 (ATFS-1), which in turn enhance mitochondrial energy metabolism and stress resilience. Our findings reveal that PLA elevates organismal ATP levels and oxygen consumption rate (OCR), which is reflected in the average movement of nematodes. Furthermore, we introduce a healthspan-associated parameter, the Healthy Aging Index (HAI), which is enhanced through PLA supplementation.

Several studies have suggested the potential mechanisms underpinning the effects of LAB on host physiology may be linked to their regulatory roles in modulating the gut microbiota ^8^. LAB are known to influence the composition of the gut microbiota, which, in turn, can regulate the production of short-chain fatty acids (SCFAs) within the gut ^9^. SCFAs, as reported in pre-clinical and clinical studies, have demonstrated positive effects on various aspects of human health, diseases, and even communication between the gut and the brain ^10,11^. In our current study, we explore the role of 3-phenyllactic acid (PLA), a major class of LAB metabolite that shares similarities with representative SCFAs such as phenylbutyric acid and phenylpropionic acid. PLA may offer the advantages associated with SCFAs, including the enhancement of stress resistance and its function as an energy substrate, resulting in a myriad of physiological processes that contribute to human health and the management of diseases ^12,13^. These observations and the effects associated with SCFAs and their production play a pivotal role in the health benefits derived from LAB supplementation.

Utilizing defined diets containing PLA into *C. elegans* experimental model allows us to further investigate the significance of probiotic diets and their impact on the aging process and other physiological functions.

## Results

### Validation of longevity-related LAB factors and their impact on *C. elegans* healthspan

In our quest to substantiate potential health advantages associated with LAB and pinpoint their effectors, we utilized LAB as a dietary component and assessed its impact on the lifespan of the *C. elegans* model. We specifically employed LAB isolated from green tea, recognized for its effectiveness as a probiotic source ^14–16^. Our findings unequivocally validate that LAB treatment significantly retards the aging process and extends the lifespan of *C. elegans* in comparison to the standard food source for these nematodes, OP50 bacteria (Figures 1A, S1A-S1B, and Table S1). Notably, a dose-response analysis of bacterial concentration indicated a substantial extension in lifespan, ranging from 6.6% to 21.2%. For subsequent experiments, we maintained a concentration of 10×10^10 bacteria ml^-1. Remarkably, LAB not only prolonged lifespan but also elicited a positive impact on various health-related factors. These encompassed a reduction in age-related lipofuscin accumulation (Figure S1C), increased average speed, enhanced coordination of body movements through increased pumping frequency (Figures S1D-S1E), and a decrease in age-related triglyceride (TG) levels (Figure S1F)^17^. Furthermore, LAB treatment conferred enhanced thermotolerance (Figure S1G) and increased resistance to acute oxidative stress (Figure S1H) without detectable alterations in the levels of major reactive oxygen species (ROS) defense enzymes, specifically superoxide dismutase (SOD) and catalase activity (Figures S1I-S1J). Importantly, in contrast to numerous other compounds associated with lifespan extension, LAB did not induce significant alterations in brood size (Figure S1K) or bacterial proliferation, reflecting calorie intake (Figure S1L). Furthermore, our observations revealed no discernible food preferences concerning food intake (Figures S2A-S2B).

**Figure 1.**
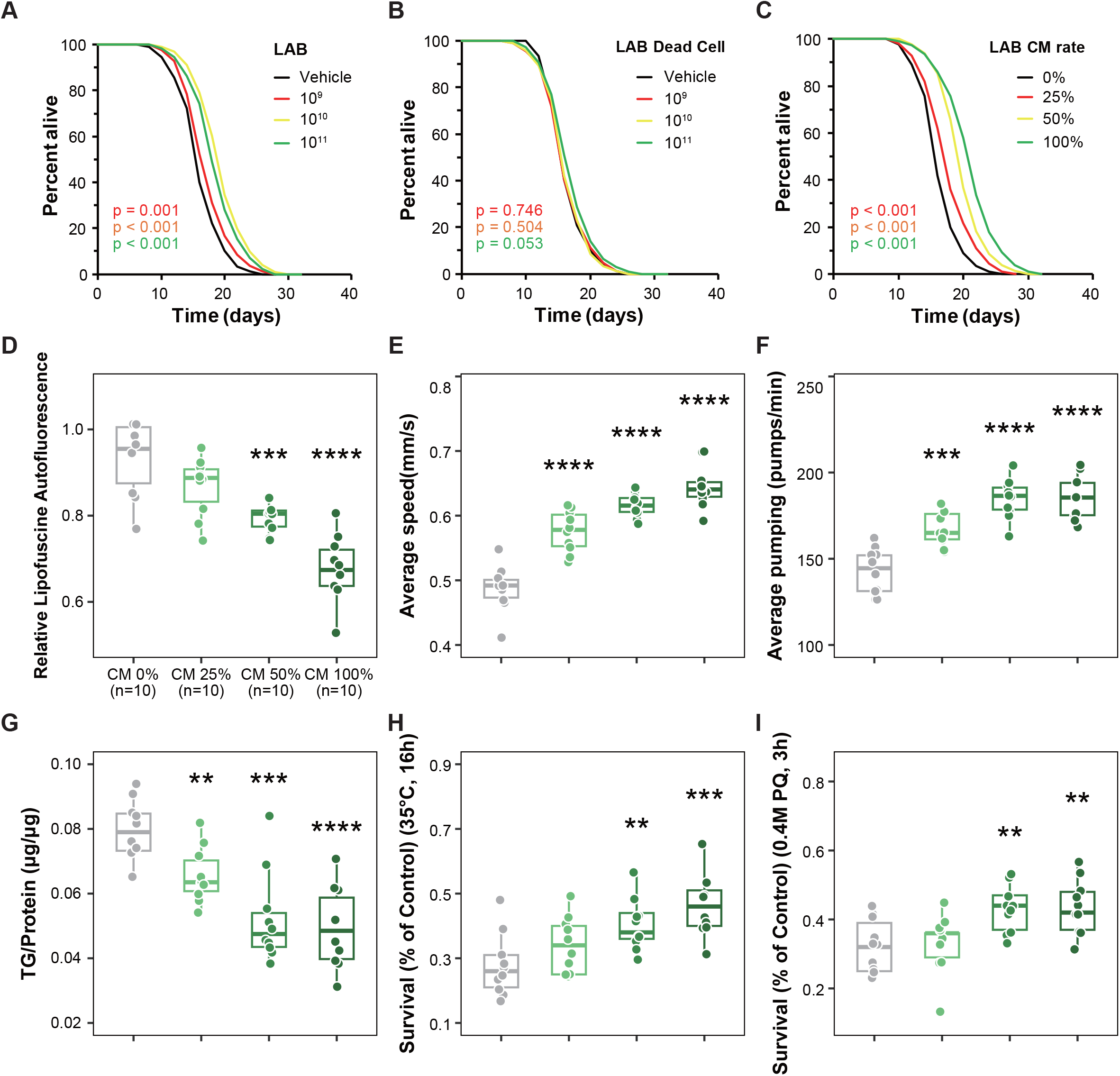
*Lactiplantibacillus* (LAB) and its conditioned media improve fitness and healthspan. (A) Effects of LAB (10 × 10^9^–10 × 10^11^ bacteria/ml) versus the vehicle (black) on lifespan (p-value listed, respectively, log-rank test); colour coding assigned to all subsequent panels. (B) Effects of LAB dead cell (10 × 10^9^–10 × 10^11^ bacteria/ml) versus the vehicle (black) on lifespan (p-value listed, respectively, log-rank test). (C) Effects of LAB conditioned media (25-100% in total media) versus the 0% CM (black) on lifespan (p-value listed, respectively, log-rank test). (D-L) Effects of CM versus the 0% vehicle in animals regarding (D) lipofuscin (***p < 0.001 and ****p < 0.0001 versus the vehicle group, one-way ANOVA, n = 20 worms × three assays each), (E) average speed (****p < 0.0001 versus the vehicle group, one-way ANOVA, n = 20 worms × three assays each), (F) average pumping (***p < 0.001 and ****p < 0.0001 versus the vehicle group, one-way ANOVA, n = 20 worms × three assays each), (G) triglyceride (TG) content (**p = 0.002, ***p < 0.001, and ****p < 0.0001 versus the vehicle group, one-way ANOVA, n =3 worm pellets), (H) thermotolerance (p = 0.169, **p = 0.008, and ***p < 0.001 versus the vehicle group, one-way ANOVA, n = 20 worms × nine measurements each), (I) oxidative stress resistance (p = 0.914, **p = 0.004, and **p = 0.002 versus the vehicle group, one-way ANOVA, n = 20 worms × nine measurements each). Lifespan assay data are represented in Table S1. Error bars represent the mean ± s.d.

To validate our hypothesis regarding the specific metabolites of LAB that enhance healthspan in organisms, we embarked on the quest to identify the characteristic metabolite responsible for explaining the favorable effects of probiotic interventions. Given that probiotics, including LAB, produce metabolites which may function as bioactive compound, we postulated that living LAB or its metabolites could impact age-associated parameters. In pursuit of this hypothesis, we designed a lifespan assay employing LAB dead cells or conditioned media (CM) derived from LAB incubation. Intriguingly, the use of heat-inactivated LAB (Figures S2C-E) or UV-killed dead cells (Figures 1B and S2F-G) at the same dose as living bacteria did not elicit alterations in the lifespan of *C. elegans*. In stark contrast, heat-inactivated OP50 bacteria noticeably extended lifespan (Table S1). This suggests that only metabolites produced by living LAB possess the capacity to influence age-related factors. Furthermore, we observed noteworthy differences in the diversity of bacterial food sources through the unweighted pair group method with arithmetic mean (UPGMA) tree and principal component analysis (PCA) using 16S metagenomic sequencing when comparing samples exposed to living LAB versus heat- or UV-treated LAB (Figure S2H-I). Of particular interest, conditioned media (CM) obtained from living LAB exerted a significant dose-dependent increase in *C. elegans* lifespan (Figures 1C, S2J-S2K, and Table S1). Varying the proportion of CM (0%, 25%, 50%, and 100%) in liquid media resulted in a dose-dependent reduction of lipofuscin (Figure 1D), an increase in average speed (Figure 1E), elevated pumping frequency (Figure 1F), reduced triglyceride content (Figure 1G), along with enhanced thermotolerance (Figure 1H) and resistance to oxidative stress (Figure 1I), similar to the effects seen with living LAB interventions (Figure S1). CM also had no discernible impact on progeny, superoxide dismutase (SOD) activity, and catalase activity, demonstrating congruence with the outcomes observed with LAB (Figures S2L-S2N). In summary, both living LAB and LAB-derived CM effectively extended healthspan, inducing thermotolerance and safeguarding against oxidative stress. These findings strongly indicate that certain metabolites contained in CM may serve as key compounds in LAB-mediated longevity.

### 3-phenyllactic acid extends age-associated health factors as an active compound of LAB

To elucidate the active constituent responsible for LAB-induced longevity, we conducted CE-TOFMS metabolome profiling on *C. elegans* fed with LAB or heat-inactivated LAB (Figure 2A). Our analysis revealed that 3-phenyllactic acid (PLA) emerged as the most prominently increased metabolite (Figure 2B), in line with previous observations^18^. As the primary product of carbohydrate fermentation by LAB, we explored whether PLA could influence the lifespan and healthspan of *C. elegans*. Based on earlier studies with lactate metabolites^19,20^ and PLA with its antimicrobial properties^21,22^, presented potential for enhancing longevity. Significantly, PLA demonstrated the most pronounced effects on lifespan extension in a dose-dependent manner (ranging from 22.7% to 23.3% at concentrations of 0.4-10 mM) (Figures 2C, S3A-B, and Table S1). In addition to increasing lifespan, supplementation with PLA also extended healthspan factors. This was evident in the suppression of lipofuscin accumulation (Figure 2D), improved mobility (Figures 2E-2F), reduced triglyceride levels (Figure 2G), and enhanced resilience to heat (Figure 2H) and oxidative stress (Figure 2I). We also considered the D-form of PLA in lifespan assays, demonstrating slightly reduced effects compared to the L-form at the same dose range for D-PLA (Figures S3C-S3E and Table S1).

**Figure 2.**
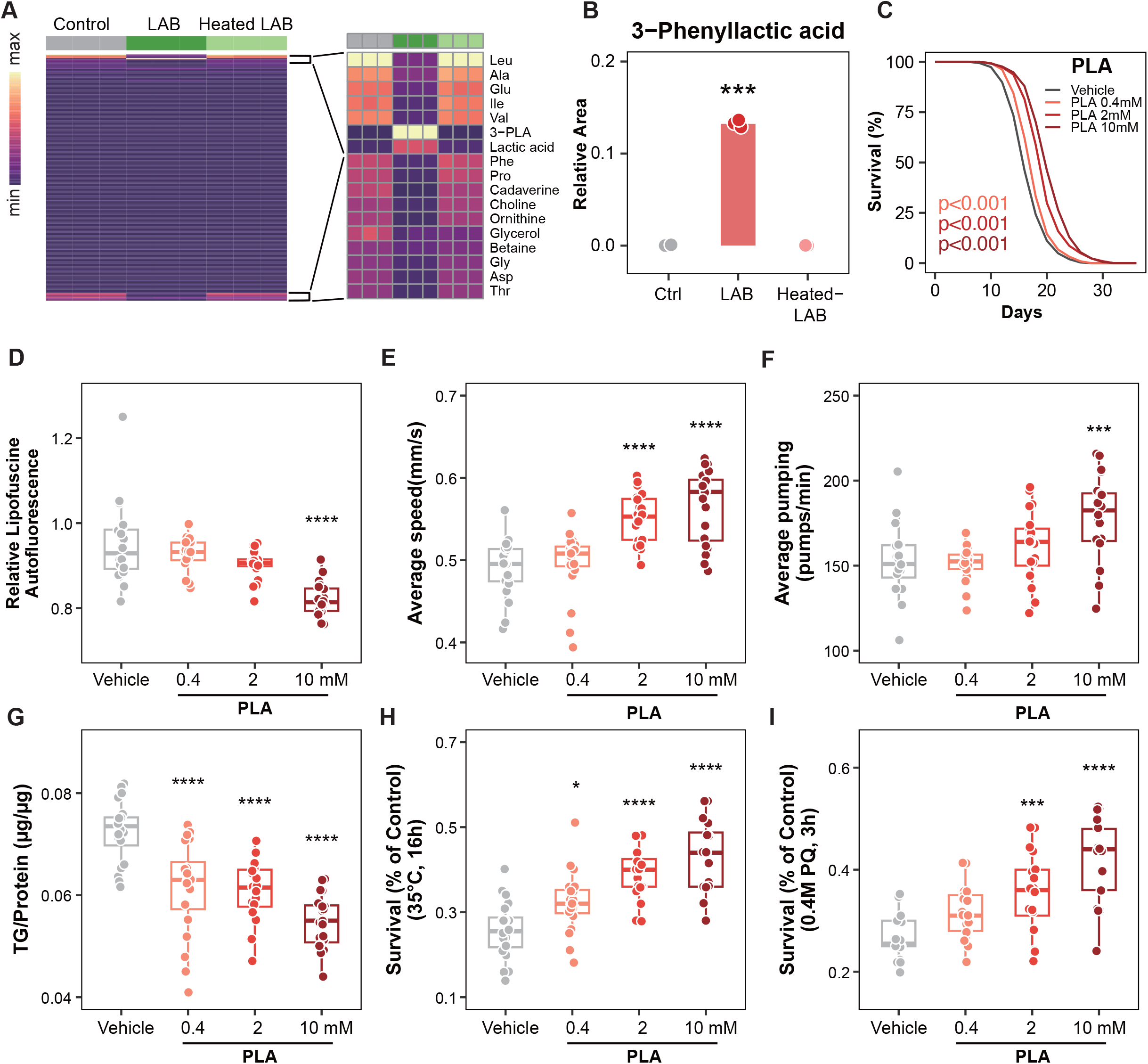
LAB-derived metabolite 3-phenyllactic acid (PLA) promotes healthy aging in *C. elegans*. (A) Metabolite profiling of the control, LAB or heat-treated LAB-fed worms using LC-TOF/MS. (B) PLA (***p < 0.001 versus the vehicle group, one-way ANOVA, n = 3 worm pellets each) was selected as increased metabolite of LAB-fed worms compared to vehicle or heat-inactivated-LAB condition (****p < 0.0001 versus the vehicle group, one-way ANOVA, n = 3 worm pellets each). (C) Effects of PLA versus vehicle on lifespan (p-value listed, respectively, log-rank test). (D-I) Effects of PLA versus control for (D) lipofuscin (****p < 0.0001 versus the control group, one-way ANOVA, n = 20 worms × three assays each), (E) average speed (****p < 0.0001, Student’s t-test, n = 10-15 worms × 3 assays each), (F) average pumping (***p < 0.001 versus the vehicle group, one-way ANOVA, n = 20 worms × three assays each), (G) triglyceride (TG) content (****p < 0.0001 Student’s t-test, n = 3 worm pellets), (H) thermotolerance (****p < 0.0001 versus the control group, one-way ANOVA, n = 20 worms × nine measurements each), and (I) oxidative stress resistance (***p < 0.001 and ****p < 0.0001 two-tailed *t*-test with FDR < 0.05, n = 20 worms × 9 measurements each). Lifespan assay data are represented in Table S1. Error bars represent the mean ± s.d.

### 3-Phenyllactic acid enhances healthy aging index and mitochondrial activity

In contemporary longevity research, the extension of healthspan holds paramount significance. It is essential to recognize that longevity does not invariably equate to healthy aging, especially in experimental settings where survival may persist despite a substantial reduction in physical activity. To facilitate a quantitative evaluation of healthy aging, we introduced the Healthy Aging Index (HAI), which encompasses three key components: (1) the assessment of physical performance through average speed, (2) evaluation of physical fitness and mitochondrial function via oxygen consumption rate (OCR), and (3) measurement of energy production and consumption efficiency using total ATP.

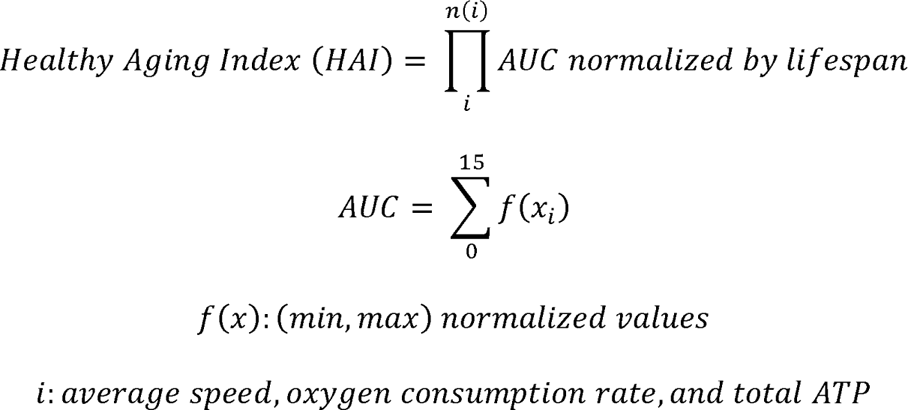

Given that PLA extends the lifespan while enhancing motility in aged worms, we sought to estimate senescence-associated health factors and introduce a novel metric for healthy aging, the Healthy Aging Index (HAI). To validate the health benefits, we conducted a comparative investigation of PLA with well-established longevity compounds such as metformin, resveratrol, LY294002, and rapamycin, using text-mining methods as described^23^. We evaluated key parameters, including average speed (Figure 3A), oxygen consumption rate (OCR) (Figure 3B), and ATP levels (Figure 3C), using the most effective doses of these compounds. The cumulative outcome was used to calculate the HAI, which represents an integrated healthy aging metric (Figure 3D). In this representation, the vehicle condition was assigned as the standard value line. Notably, PLA exhibited the highest HAI value, followed by metformin and resveratrol, while LY294002 and rapamycin displayed HAI values lower than the standard value. As anticipated, it became apparent that HAI did not align perfectly with lifespan (Figure 3E). The equation used to determine HAI underscores that it encapsulates healthspan, going beyond simple lifespan assessment (Figure 3F). These findings collectively suggest that PLA is a potent inducer of healthy aging, with a particular focus on enhancing organismal motility and energy metabolism.

**Figure 3.**
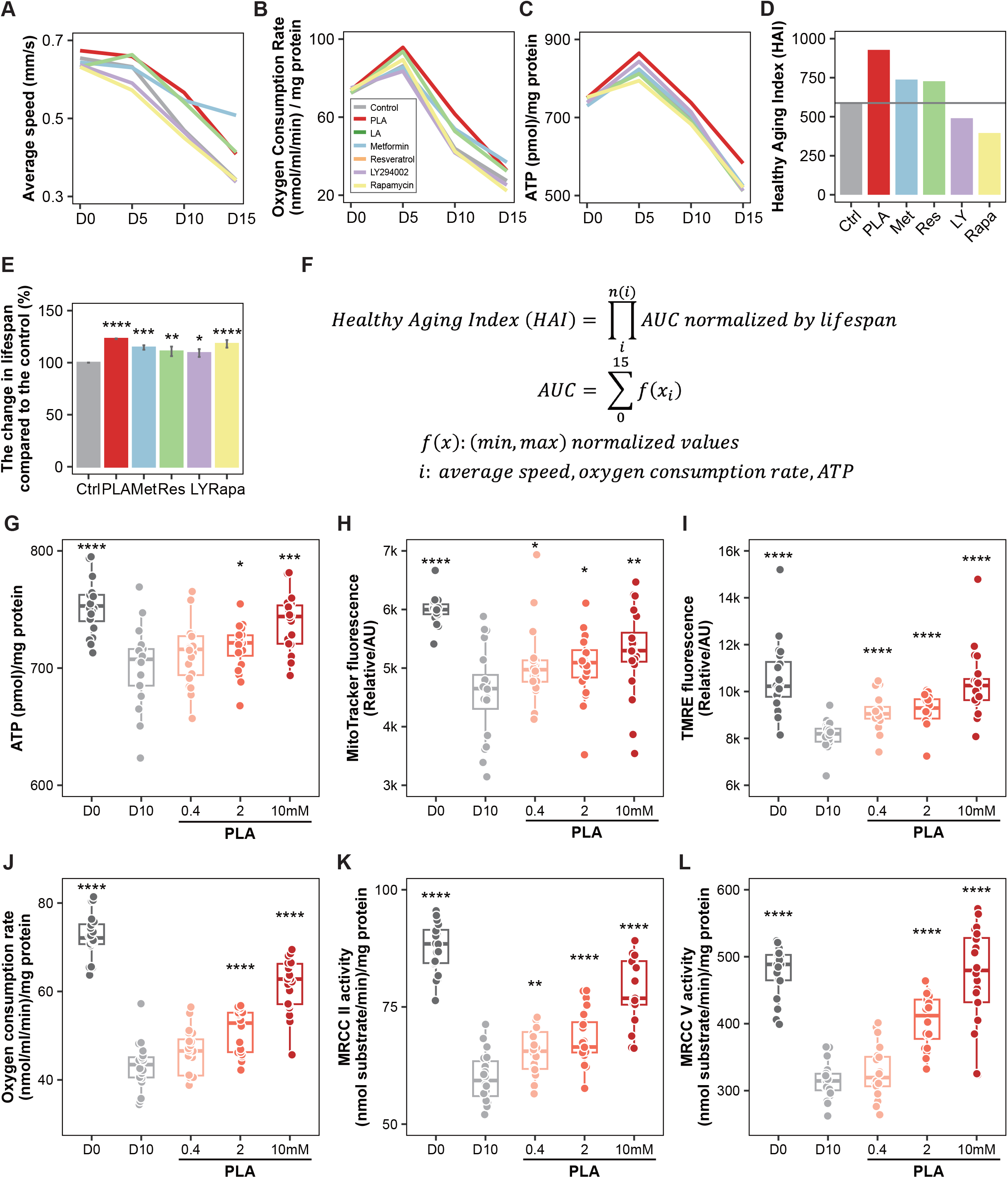
PLA supplementation enhances HAI and mitochondrial oxidative phosphorylation complex activity. Age-related comparison of longevity compound PLA, Metformin, Resveratrol, LY294002, or Rapamycin for (A) average speed, (B) oxygen consumption rate (OCR), and (C) ATP contents. (D) Age-associated HAI and (E) lifespan changes. Healthy Aging Index (HAI) was estimated by the (F) following equation. (G) Organismal ATP level under PLA treatment (***p < 0.001 and ****p < 0.0001 versus control worms, n =3 worm pellets). (H-I) Mitochondrial activity was examined by (H) mitochondrial mass with MitoTracker green (*p < 0.05, **p < 0.01, and ****p < 0.0001 versus the control group, one-way ANOVA, n = 20 worms × nine measurements each) and (I) mitochondrial membrane potential with TMRE (****p < 0.0001 versus the control group, one-way ANOVA, n = 20 worms × nine measurements each). (J) The oxygen consumption rate examined in PLA treated condition (****p < 0.0001 versus control worms, n =3 worm pellets). (K-L) The enzyme activity of mitochondrial respiratory chain complex (MRCC) (K) II (**p = 0.018 and ****p < 0.0001 versus control worms, n =3 worm pellets) or (L) V was evaluated in worms (****p < 0.0001 versus control worms, n =3 worm pellets). Error bars represent the mean ± s.d.

Until now, lactate has been primarily regarded as a metabolic waste product in energy metabolism, differing from glucose, which is traditionally recognized as a fuel source. However, emerging evidence in mammalian systems has shed light on the potential of lactate as a significant circulating carbohydrate fuel source, as well as its role in equilibrating the NADH/NAD ratio alongside pyruvate^24–27^. In this context, we find a noteworthy example in the metabolic myopathy McArdle disease, where a depletion of lactate can induce exercise intolerance and muscle fatigue^28^. Of particular importance is the role of lactate in ATP generation, a primary energy carrier in all cellular processes. We observed that, especially during the aging stages (from Day 5 to Day 15), ATP production increased significantly with PLA interventions. Furthermore, the amount of ATP was notably elevated in PLA-fed worms (Figure 3G). Given the potential harm of excessive acid accumulation, exemplified by conditions such as lactic acidosis^29,30^, we assessed the impact of high doses of PLA on longevity in a dose range of 25-100 mM. Intriguingly, PLA not only failed to extend lifespan but, more importantly, did not reduce lifespan when administered at high doses (Figures S3F-S3H and Table S1). In conclusion, metabolites derived from LAB, namely PLA, extend the healthspan of *C. elegans* independently. PLA exhibits a safe and significant lifespan-extending effect, primarily driven by improved health factors, with little or no observed side effects.

Given the profound implications of energy generation on age-related mitochondrial disorders and mitochondrial activities, we proceeded to validate the decline in mitochondrial function associated with aging. In Day-10 worms treated with PLA, we observed a significant decrease in mitochondrial function, as evidenced by reduced levels of generated ATP (Figure 3G), decreased mitochondrial volume (Figure 3H), compromised membrane potential (Figure 3I), and diminished oxygen consumption rate (OCR) (Figure 3J). We recognized that mitochondrial activity is intricately linked to the potency and efficiency of mitochondrial respiration chain complexes (MRCC)^31^. Thus, we examined the activities of the five mitochondrial enzymatic complexes under PLA intervention. Interestingly, PLA specifically interacted with MRCC II (Figure 3K) and MRCC V (Figure 3L), while no interactions were observed with the remaining MRCCs (Figures S4A-S4C). This interaction was notable for its specificity and surprising similarity to the effects of lactate administration on MRCC in mouse skeletal muscle^32^. Furthermore, in accordance with previous studies on bioenergetic health assessment in *C. elegans*^33,34^, we observed a dose-dependent increase in oxygen consumption rate under PLA treatment. The observed higher ATP generation and MRCC activity in worms suggested that PLA-mediated longevity may involve a context similar to that initiated by stress stimulation. This increase in ATP induced the extension of healthspan, further supporting the idea that PLA enhances longevity, rather than LAB itself.

### SKN-1 and ATFS-1 mediates PLA-dependent longevity

In our pursuit of unraveling the molecular mechanisms underlying the longevity effects of PLA, we conducted RNA sequencing (RNA-seq) on nematodes that had been fed PLA for a span of 5 days. This analysis unveiled 2,285 upregulated and 1,780 downregulated differentially expressed genes (DEGs) with a |Fold change| of ≥ 2 and p-value < 0.05, in comparison to control worms that were fed the vehicle alone. These marked alterations in the transcriptome indicated a significant metabolic shift and changes in senescence-regulatory pathways. Amidst the pool of DEGs, we focused our attention on 6 upregulated and 16 downregulated genes that exhibited particularly dramatic changes (Table S2). Given the pivotal role of these specific DEGs in longevity, we further explored their interactions with mitochondrial activation and stress resistance. Utilizing the RNA-seq data of PLA-fed worms and referencing published RNA-seq data of *skn-1* RNAi-treated worms or *cco-1* and *atfs-1* RNAi-treated worms, we were able to confirm that the majority of putative PLA-target genes were under the influence of these two prominent stress-induced transcription factors, *skn-1* (1,824 genes) and *atfs-1* (1,992 genes). Remarkably, 11.3% of genes were found to be regulated by PLA treatment, *skn-1*, and *atfs-1* activation simultaneously (Figure 4A). Based on this analysis, we propose a hypothesis that PLA instigates *skn-1*/*atfs-1*-mediated mitochondrial activation and enhances stress resilience, ultimately leading to an extension of healthspan (Figure 4B).

**Figure 4.**
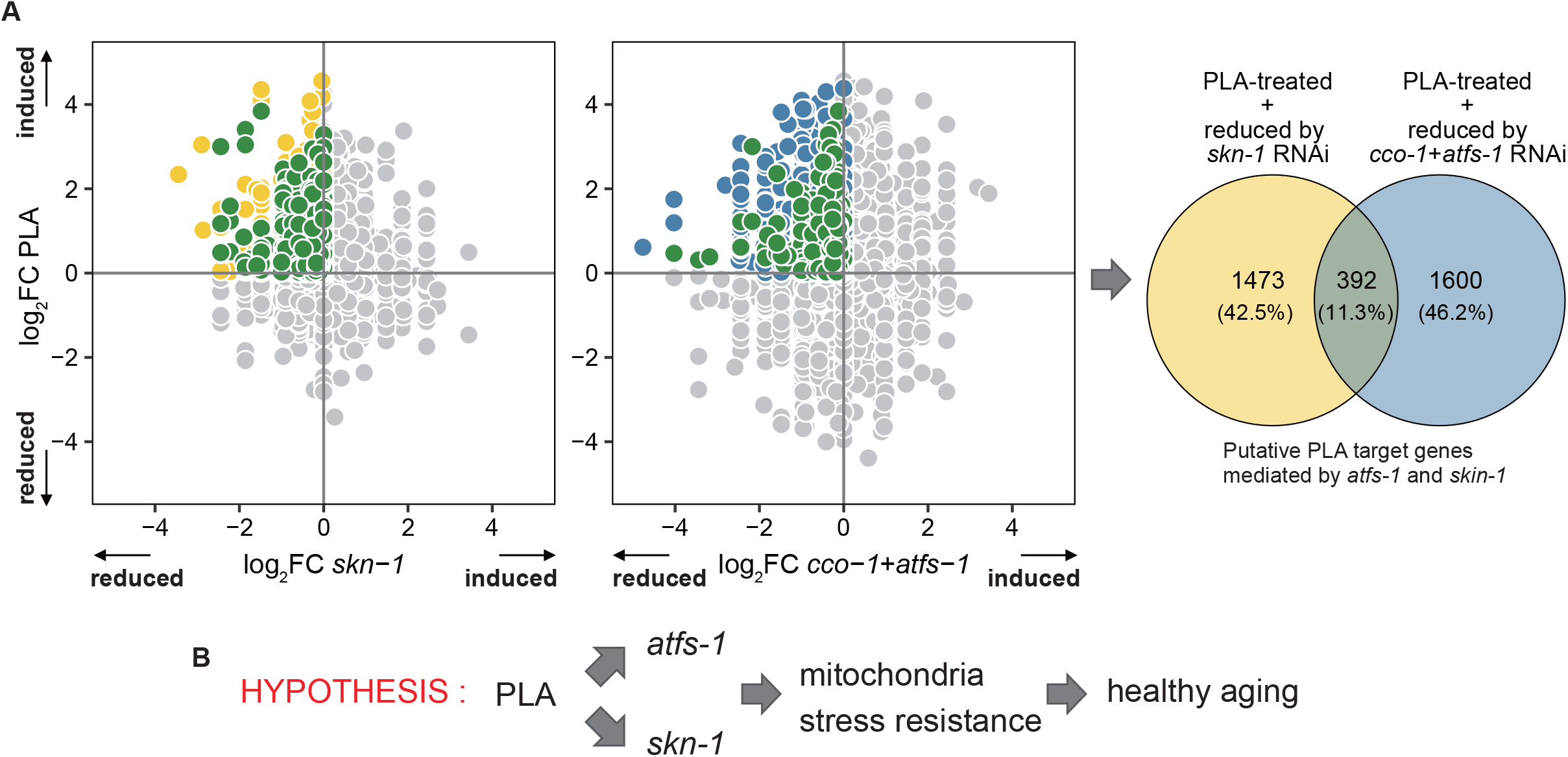
PLA supplementation activate stress-response transcription factors SKN-1 and ATFS-1. (A) Comparing the transcriptomes of the PLA-treated worm with those treated with either skn-1 RNAi or atfs-1 RNAi via scatter plots and Venn diagrams suggests that SKN-1 and AFTS-1 are crucial players generating the effect of PLA supplementation at the transcriptional level. (B) Hypothesis of PLA-induced longevity. PLA elevates organismal HAI with SKN-1/ATFS-1-mediated mitochondrial activation and stress resilience.

To test this hypothesis, we performed lifespan assay, nuclear translocation analysis, and stress resistance assay with loss of SKN-1 or ATFS-1 mutants, *skn-1*(zj15) and *atfs-1*(*gk3094*) to assess the relation with PLA effects (Figure 5). In both mutants, PLA failed to extend the lifespan, strongly suggest that PLA prolongs the *C. elegans* lifespan SKN-1 and ATFS-1-dependent manners (Figures 5A and 5D). Then, we examined whether the nuclear translocation of the transcription factors SKN-1 and ATFS-1 shifted in response to PLA supplementation. Our findings revealed that PLA induced the nuclear translocation of SKN-1 (ranging from 26.9% to 75.4% at concentrations of 0.4-10 mM) (Figures 5B-5C) and ATFS-1 (ranging from 29.7% to 60.3% at concentrations of 0.4-10 mM) (Figures 5E-5F) in a dose-dependent manner. Drawing from previous research on the mitochondrial import efficiency of SKN-1 and ATFS-1, which correlates with the degree of mitochondrial dysfunction and exposure to external stressors ^35,36^, we found that the increased nuclear import of SKN-1 and ATFS-1 coincided with consistent ATP production under PLA treatment. Notably, this effect was not altered in deletion mutants of *atfs-1*(*gk3094*) or *skn-1*(*zj15*) (Figure 5G). Additionally, thermotolerance remained unaltered in the absence of these genes when subjected to PLA treatment (Figure 5H). Furthermore, the oxidative stress resistance of *atfs-1*(*gk3094*) or *skn-1*(*zj15*) mutants in conjunction with PLA treatment did not exhibit significant changes (Figure 5I). These collective results support the hypothesis that increasing PLA interventions can mitigate compromised stress resistance by activating SKN-1/ATFS-1 and associated mitochondrial activation.

**Figure 5.**
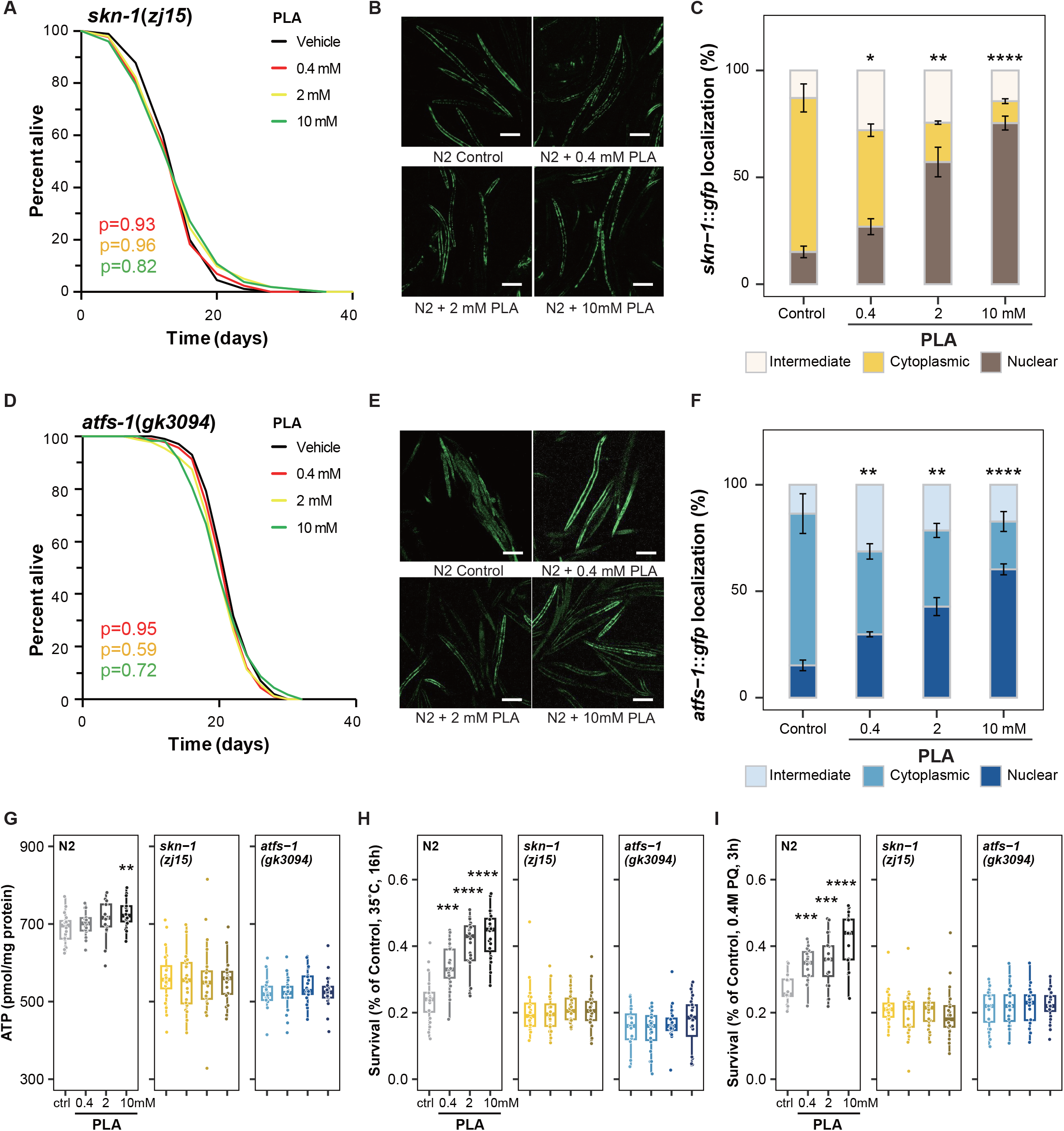
PLA mediates SKN-1 and ATFS-1 pathway. (A) Lifespan analysis of *skn-1*(*zj15*) with PLA treatment was performed. (B) Nuclear translocation of SKN-1::GFP was visualized using *skn-1::gfp* worms with PLA supplementation. Scale bars, 200 μm. (C) The quantification of SKN-1::GFP localization with PLA treatment (*p < 0.05 and ***p < 0.001 versus the control group, n = 20 worms x six measurements each). (D) Lifespan assay of *atfs-1*(*gk3094*) under PLA treatment. (E) ATFS-1::GFP localization was depicted with *atfs-1::gfp* worms under PLA treatment. Scale bars, 200 mm. (F) ATFS-1::GFP localization was quantified (*p < 0.05, **p < 0.01, and ***p < 0.001 versus the control group, n = 20 worms × six measurements each). (G) Organismal ATP level with PLA was estimated in *atfs-1*(*gk3094*) or *skn-1*(*zj15*) deletion mutant (**p < 0.01 versus the control group, n = 20 worms × six measurements each). (H) Thermotolerance assay of N2, *atfs-1*(*gk3094*) or *skn-1*(*zj15*) mutant worms under treatment of PLA (***p < 0.001 and ****p < 0.0001 versus the control group, n = 20 worms × nine measurements each, respectively). (I) Oxidative stress resistance of N2, *atfs-1*(*gk3094*) or *skn-1*(*zj15*) mutant animals with PLA supplementation (***p < 0.001 and ****p < 0.0001 versus the control group, n = 20 worms × 9 measurements each). Overall differences between conditions were analysed by two-way ANOVA. Differences in individual values or between two groups were determined using two-tailed *t*-tests (95% confidence interval). Error bars represent the mean ± s.d.

### The correlation between PLA and human muscle activity and the healthspan pathway

Healthy aging, as defined by the World Health Organization, refers to the process of developing and maintaining functional abilities that support well-being in older age ^37^. In the context of humans, healthy aging is closely associated with conditions such as sarcopenia, frailty, and cognitive impairment. To explore the potential connection between blood PLA levels and physical performance in humans, we collected plasma samples from 24 patients diagnosed with sarcopenia and 25 age-matched individuals without sarcopenia. Subsequently, we conducted relative PLA quantification using liquid chromatography–mass spectrometry (Figure 6A). The results indicated that patients with sarcopenia had significantly lower levels of PLA in their blood, and they exhibited poorer physical performance metrics, including grip strength, gait speed, the 5-time sit-to-stand test, and the short physical performance battery (Figure 6B). These findings suggest that PLA may play a role in contributing to sarcopenia and warrant further investigation through both animal and human studies.

**Figure 6.**
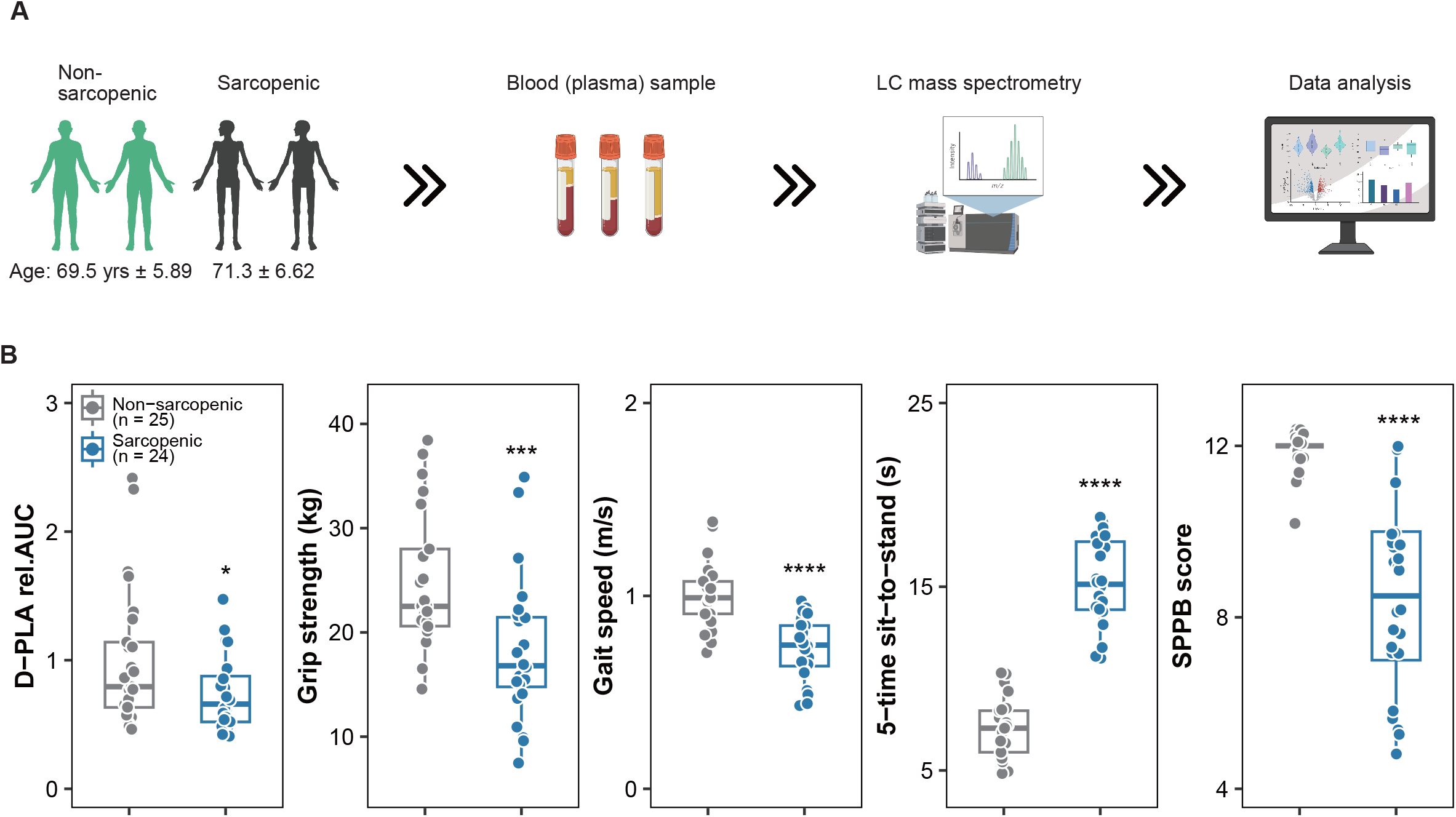
The correlation of blood PLA concentration and physical performance with human patients with sarcopenic phenotype. (A) Non-sarcopenic (69.5 ± 5.89 years) or patients with sarcopnic phenotype (71.3 ± 6.62 years) blood samples were collected and D-PLA concentration in human blood samples was measured. Muscle mass and Skeletal muscle mass index of non-sarcopenic or sarcopenic participants. Muscle function of participants were evaluated with Grip strength (****p < 0.0001), Gait speed (****p < 0.0001), FTSS (****p < 0.0001), and SPPB score (****p < 0.0001).

## Discussion

In our pursuit of identifying commonly increased metabolites that might impact longevity in LAB-mediated long-lived organisms through an organismal approach with *C. elegans*, we examined the effects of live bacteria, conditioned media, and dead LAB cells. LABs have gained industrial importance and are generally recognized as safe (GRAS) due to their widespread presence in food and their host health benefits, including those to human mucosal surfaces ^38,39^. Among the LAB metabolites, our focus was on 3-phenyllactic acid (PLA), the most abundantly produced metabolite ^15,40^. Notably, previous studies indicated that with aging, the proportion of lactate decreased in *C. elegans* models, and the expression of lactate dehydrogenase played a role in regulating longevity and neurodegeneration in *Drosophila* models ^19,41,42^. Despite the long-held perception of lactate as a metabolic waste product associated with excessive respiration and skeletal muscle fatigue, or even detrimental effects leading to lactic acidosis ^30,43^, recent research has shifted our perspective on lactate. It is now recognized as a key player in organismal maintenance processes, serving as a major energy source for mitochondrial respiration, a primary gluconeogenic precursor that safeguards muscle contractions, and a crucial signaling molecule ^24,44^. Estimates of circulating metabolite flux in mice have revealed that lactate exists in every tissue of the body as intermediates of the tricarboxylic acid (TCA) cycle ^27^. Furthermore, recent in vivo studies comparing lactate and glucose metabolism have shown that lactate predominates in feeding the TCA cycle, even in tumors ^45^. Notably, given its safety as a common food additive, lactate supplementation has demonstrated various clinical health benefits, including the mitigation of ethanol-induced gastric mucosal damage, the prevention of inflammatory bowel disease, and the development of intestinal epithelium as mucosal delivery vehicles ^46–48^. These studies collectively highlight the potential impact of lactate and its metabolites on energy metabolism and healthspan extension at the organismal level. Circulating lactate facilitates the decoupling of carbohydrate-derived mitochondrial energy metabolism from glycolysis, and the lactate/pyruvate ratio helps balance the NADH/NAD^+^ ratio across cells and tissues ^49^. This mechanism supports mitochondrial energy generation as a carbohydrate fuel source, shedding light on its positive influence on aging^24,44,50^.

Encouraged by these findings, we delved deeper into the potential advantages of PLA for organismal fitness and mitochondrial activity. Guided by the perspectives on the roles of energy metabolism, we explored phenylpyruvic acid, a metabolic precursor of PLA, which was predicted as an NAD^+^ activator through in silico analysis ^51^. The observed increase in longevity was found to be contingent on its conversion to PLA. Notably, the elevated energy generation achieved through PLA interventions further enhanced mitochondrial activity, contributing positively to overall vitality. Within the context of stress resilience, it’s important to recognize that external stress accumulates over the adult lifetime of *C. elegans* ^52^, resulting in the presence of abundant reactive oxygen species that can inflict substantial damage on proteins and lipids, ultimately leading to the onset of metabolic diseases and a shortened lifespan ^53,54^. Enhancing tolerance to external stress factors has been previously linked to the promotion of healthspan in *C. elegans* and other model organisms ^55,56^. In our study, we directed our attention to investigating the actions of SKN-1 and ATFS-1, two transcription factors identified through transcriptome analysis of PLA-fed animals, in the context of stress resistance. In our current study, the CM of LAB, as well as PLA, significantly increased thermotolerance and resistance to acute oxidative stress. Notably, these stress tolerance effects were entirely abolished in the absence of the SKN-1 and ATFS-1 transcription factors. Furthermore, our study provided evidence that PLA upregulates stress resilience through a mechanism involving the well-known stress-related transcription factors SKN-1/NRF2^57–59^ and ATFS-1^36,60^, subsequently enhancing the hallmark of stress resilience, namely, survival under conditions of thermotolerance and oxidative stress^61^.

In our study, we propose an alternative perspective on how PLA achieves lifespan and health benefits in *C. elegans* by addressing the toxicity associated with the dark side of lactate accumulation. Furthermore, considering that lactate accounts for approximately 25% of muscle content, we explored the relevance of PLA to age-related muscle function. In our nematode model, PLA effectively ameliorated age-associated motility decline, as evidenced by improvements in movement speed and pumping rate. This led us to investigate age-associated muscle atrophy, specifically sarcopenia. Notably, a clear inverse relationship between the concentration of PLA in sarcopenia patients and reduced muscle function was observed. It is intriguing to observe the potential connection between a specific metabolic compound and human physical performance. While our work requires additional studies to determine whether the effects on fitness and longevity can be translated to humans, this study provides a valuable clue that PLA has the potential to activate cellular energy metabolism and enhance stress resilience, either through LAB-derived supplementation or as a standalone intervention. This offers a promising avenue for promoting organismal vitality.

## STAR METHODS

Detailed methods are provided in the online version of this paper and include the following:

- KEY RESOURCES TABLE
- CONTACT FOR REAGENT AND RESOURCE SHARING
- EXPERIMENTAL MODEL AND SUBJECT DETAILS

◦ Nematode strains and maintenance
◦ Worn synchronization and food preparation
- METHOD DETAILS

◦ LAB isolation and growth condition
◦ Lifespan analysis
◦ Lipofuscin analysis
◦ Movement, pharyngeal pumping rates, and fertility assay
◦ Triglyceride quantification
◦ Stress resistance assays
◦ Superoxide dismutase and catalase activity assay
◦ Preparation of chemical supplementary plates
◦ RNA sequencing, data analysis, and visualization
◦ RNAi in *C. elegans*
◦ RNA extraction and quantitative real-time PCR
◦ Assay for MGO and glyoxalase activity
◦ Component levels of lactic and phenyllactic acid metabolic pathway
◦ Mitochondria preparation and measurement of MRCC activity
◦ Measurement of ATP generation
◦ SKN-1::GFP, ATFS-1::GFP, and HSF-1::GFP localization assay
◦ Quantification of ferroptosis level
◦ Estimation of PLA amount in sarcopenic blood sample
◦ Spearman and Pearson correlation method
- STATISTICAL ANALYSIS
- DATA AVAILABILITY

## SUPPLEMENTAL INFORMATION

Supplemental information includes seven figures, two tables, and can be found with this article online

## ACKNOWLEDGEMENTS

JK, YJ^1^ and DR were supported by the National Research Foundation of Korea (NRF) grant funded by the Korea government (MSIT) (2022R1I1A1A01063460, 2023R1A2C3006220, and 2022K2A9A1A06091879) and by the GIST Research Institute (GRI) IIBR grant funded by the GIST in 2023. HSY was supported by a grant from the Korea Health Technology R&D Project, through the Korea Health Industry Development Institute (KHIDI), funded by the Ministry of Health & Welfare, Republic of Korea (HR22C1734). *C. elegans* strains used in this research were provided by the *Caenorhabditis* Genetics Center (University of Minnesota, USA), which is funded by the NIH Office of Research Infrastructure Programs.

## AUTHOR CONTRIBUTIONS

The study was conceptualized and designed by JK, CD, and DR. JK was responsible for conducting the majority of the experiments. YJ^1^ handled the analysis and visualization of all experimental data, which included all raw data, transcriptomic and metabolomic data and sarcopenic blood data. LAB production and metabolomics of the LAB CM, which led to the identification of PLA, were carried out by YJ^3^, JHR, WGK, and DC. HSY was in charge of conducting human studies and collecting blood samples. PLA measurements in human plasma using LC-MS were performed by Gland DWC. The manuscript was written by JK, DC, and DR; with DR also handling the editing. The research was conceived, designed, analyzed, interpreted, and overseen by JK and DR.

## STAR★METHODS

### KEY RESOURCES TABLE

#### CONTACT FOR REAGENT AND RESOURCE SHARING

Further information and requests for resources and reagents should be directed to and will be fulfilled by the Lead Contact, Juewon Kim and Dongryeol Ryu.

#### EXPERIMENTAL MODEL AND SUBJECT DETAILS

##### Nematode strains and maintenance

*C. elegans* strains were maintained on Nematode Growth Media (NGM) agar plates at 20°C using OP50 bacteria as a food source. The following strains were used (strain, genotype): Bristol N2, wild type; AY135, *zcIs4 V*; QV225, *skn-1*(*zj15*); LG333, *skn-1*(*zu135*)IV; gels7; VC3201, *atfs-1(gk3094)V*; OP675, *unc-119*(*tm4063*) III; wgls675. They were all obtained from the Caenorhabditis Genetics Center (CGC). All strains are listed in Key Resource Table.

##### Worm Synchronization and food preparation

To obtain synchronized nematode, six unmated hermaphrodites were placed on a nematode growth medium (NGM) plate with sufficient food. After 4 days, lysed F1 progenies that were selfing hermaphrodites yielded synchronized hermaphrodite eggs. The obtained eggs were grown on NGM plates. OP50 bacteria or lactic acid bacteria (LAB) were cultured at 37 °C in LB or MRS medium for up to 12 hours. The bacteria concentration was calculated by a combination of counting the colonies and measuring the OD at a wavelength of 600 nm. The OP50 was centrifuged, resuspended and diluted in H_2_O at bacteria concentrations of 10×10^11^ bacteria ml^-1^ or 10×10^9^, 10×10^10^, or 10×10^11^ for the LAB and LAB dead cell experiments until day 4 of feeding beginning at day 1 of adulthood. The conditioned media of LAB were mixed with M9 buffer to 25%, 50%, or 100% in liquid culture environment.

### METHOD DETAILS

#### LAB isolation and growth condition

*Lactiplantibacillus* APSulloc 331261, isolated from green tea (Dolsonfield, Jeju island, Korea) and its isolation, characterization, safety, and biological properties were described previously ^14–16^. The designation of this strain has been amended with its trademark (GTB1^TM^) and further reference to this strain will be as lactic acid bacteria (LAB). LAB was cultured at 37°C for 18 h and prepared on a daily basis in MRS broth (Difco Laboratories, NJ, USA). To preparation of cell-free conditioned media (CM), overnight culture of LAB was sub-cultured to mid-to-late logarithmic phase and adjusted to OD_600_ = 1.0. LAB-derived supernatant was acquired after two centrifugation steps at 15,000 × *g* for 10 min. Then, supernatant was filtered using 0.45 μm and 0.2 μm syringe filters (Whatman, Maid-stone, UK) and stored at -20°C until use. CM (100%) was diluted to concentrations of 50% and 25% (v/v) with S-basal buffer and cultured in liquid media ^62^. The heat-inactivation of LAB was obtained by putting culture suspension in 70°C water bath for 40 min with periodical mixing and LAB dead cell was prepared with two freeze thaw cycles using liquid nitrogen and a 70°C water bath.

#### Lifespan analysis

Lifespan assays were carried out at 20°C on NGM agar plates using standard protocols and were replicated in three independent experiments. *C.elegans* were synchronized with 70 μl M9 buffer, 25 μl bleach (10% sodium hypochlorite solution) and 5 μl of 10N NaOH. Either a timed 64 hours of egg lay or egg preparation, around 150 young adults were transferred to fresh NGM plates. All compounds were blended into the NGM media after autoclaving and before solidification except condition of liquid media. Living worms were transferred to assay plates, which were seeded with OP50 or a designated RNAi feeding clone with 50 μg ml^-1^ ampicillin (Sigma-Aldrich, MO, USA) every other day. Worms that crawled of the plates or exhibited internal progeny hatching were censored and showed no reaction to gentle stimulation were scored as dead. Lifespan data were analyzed using R software (ver. 4.1.0, “coin” package); Kaplan-Meire survival curves were depicted, and *P* values were calculated using the log-rank (Mantel-Cox) test. All lifespan data are described in Table S1.

#### Lipofuscin (age pigment) analysis

Nematode were synchronized and treated for 10 days with vehicle, LAB or chemicals from L4 larvae stage (Day 0). For measuring lipofuscin accumulation, Day 10 worms washed with M9 buffer 3 times and distributed on a 96-well plate (Porvair Sciences black with glass-bottomed imaging plates, #324002). Lipofuscin auto-fluorescence was determined using a fluorescence plate reader (Synergy H1, BioTek, VT, USA; excitation: 390-410, emission: 460-480) with normalized to the stable signal of the worms (excitation: 280-300, emission: 320-340) as blank.

#### Movement, pharyngeal pumping rates, and fertility assay

Day 10 of 10 – 15 worms were transferred to fresh NGM plates and recorded 30 second movement with a microscope system (Olympus SZ61 microscope with Olympus camera eXcope T300, Olympus, Tokyo, Japan). Subsequently, 5 independent movement clips per experimental condition were analyzed with TSView 7 software (ver. 7.1) and average speed was calculated as distance (mm) per second. The pharyngeal pumping of 30 worms per each condition were assessed for 1 min using a SZ61 microscope (Olympus, Tokyo, Japan). Pharyngeal contractions were recorded with Olympus camera eXcope T300 at 18-fold optical zoom and videos were played back at 0.5× speed and pharyngeal pumps counted. For fertility assay, worms were synchronized and single L4 nematode were transferred on single plates applying vehicle or LAB or reagent then moved to fresh plates. Progeny of worms were allowed to hatch and counted.

#### Triglyceride (TG) quantification

TG contents were measured using the triglyceride colorimetric assay kit (Abcam, MA, USA) with manufacturer’s protocol. Briefly, frozen worm pellets in liquid nitrogen with 5% Triton X-100. The pellets were sonicated and diluted for protein determination by BCA assay (Pierce, IL, USA). The samples were heated till 80°C and shaken for 5 min then, samples cool-downed to room temperature to solubilize all the TG. TG contents were normalized relative to protein contents and three independently collected worm pellets were assayed for each experimental condition.

#### Stress resistance assays

Synchronized worms treated each condition for 10 days. For the oxidative stress assay, 20 worms were placed on solid NGM plates containing 0.4M paraquat (PESTANAL, Sigma-Aldrich, MO, USA) for 3 hours. Assays repeated for 9 times and the paraquat plates were freshly prepared on the day of the assay. For the thermotolerance assay, 18-28 worms were exposed 35°C for 16 hours and survival worms were counted. Assays performed triplicate and survival rate was estimated.

#### Superoxide dismutase and catalase activity assay

To assess antioxidative enzyme activities, 10-day-old worms grounded in liquid nitrogen. A superoxide dismutase (SOD) and catalase activity levels were measured with the SOD colormetric activity kit (Invitrogen, CA, USA) or Catalase activity colorimetric/fluorometric assay kit (Biovision, CA, USA), respectively. Worm pellet samples were assessed according to manufacturer’s protocol in triplicate. SOD and catalase activity was determined using standard curves followed by normalization to protein concentration using the BCA protein assay (Pierce, IL, USA).

#### Preparation of chemical supplementary plates

The L- or D-phenyllactic acid was acquired (Sigma-Aldrich, MO, USA) and stock of 1 M was made by dissolving in water. The pH of stock solution was adjusted to 6.0 - 6.5, the same pH as the vehicle plates, with sodium hydroxide and chemical stock solution sterilized with 0.45 μm Millipore filter (Millipore, MA, USA) before use. Chemical treatment was performed by supplementing the compound in heat-inactivated OP50 (70°C, 40 min with 1M magnesium sulfate and 5 mg/ml cholesterol), unless it applied with RNAi clones.

#### RNA sequencing (RNA-seq), data analysis, and visualization

RNA was extracted and assigned using Bioanalyzer 2100 (Agilent Technologies, CA, USA) in combination with RNA 6000 nano kit. Matched samples with high RNA integrity number scores were proceed to sequencing. For library preparation an amount of 2 mg of total RNA per sample was processed using Illumina’s TruSeq RNA Sample Prep Kit (Illumina, CA, USA) following the manufacturer’s instruction. Quality and quantity of the libraries was determined using Agilent Bioanalyzer 2100 in combination with FastQC v0.11.7, Sequencing was done on a HiSeq4000 in SR/50 bp/high output mode at the Macrogen Bioinformatics Center (Macrogen, Seoul, Korea). Libraries were multiplexed in five per lane. Sequencing ends up with 35 Mio reads per sample. Sequence data were extracted in FastQ format using bcl2fastq v1.8.4 (Illumina) and used for mapping approach. FASTQ output files were aligned to the WBcel235 (February 2014) *C. elegans* reference genome using STAR^63^. These files have been deposited at the Sequence Read Archive (SRA) with the accession number PRJNA827000 and PRJNA894441. Samples averaged 75% mapping of sequence reads to the reference genome. We filtered out transcripts with Trimmomatic 0.38 platform ^64^. Differential expression analysis was performed using HISAT2 version 2.1.0, Bowtie2 2.3.4.1, with StringTie version 2.1.3b ^65,66^. Human ortholog matching was performed using WormBase, Ensembl, and OrthoList2 ^67^. Gene lists were evaluated for functional classification and statistical overrepresentation with Database for Annotation, Visualization, and Integrated Discovery (DAVID) version 6.8 and Kyoto Encyclopedia of Genes and Genomes (KEGG) pathway database.

#### RNAi in C. elegans

For RNAi-mediated gene knockdown, we fed worms HT115 (DE3) expressing target gene double stranded RNA (dsRNA) from the pL4440 vector. The clones for RNAi against genes were obtained from the *C. elegans* ORF-RNAi Library (Thermo Scientific, Horizon Discovery, MA, USA). All clones were verified by sequencing (Macrogen, Seoul, Korea) and efficient knockdown was confirmed by quantitative RT-PCR (qRT-PCR) of the mRNA. Bacteria were spotted on NGM plates containing additionally 1 mM isoprophyl-β-D-thiogalactoside (IPTG) (Sigma-Alrich, MO, USA). Incubation with dsRNA initiated 64 hours after population synchronization and then young adult worms transferring to the respective treatment plates. We confirmed the RNAi knockdown of *atfs-1* and *skn-1* by qRT-PCR and transcripts levels was reduced by 74% and 87%, respectively in larvae that were cultivated on corresponding dsRNA-expressed bacteria.

#### Mitochondria preparation and measurement of MRCC activity

Mitochondria were isolated according to previously described method. Briefly, approximately 1 g of age-synchronized worms were harvested from the cultures, cleaned and washed with M9 buffer and suspended in mitochondrial isolation buffer (Abcam, MA, USA). The worm suspension was ruptured using a TissueLyser II (Qiagen, Hilden, Germany) high speed bench-top homogenizer and an equal volume of mitochondrial isolation buffer containing 0.4% BSA was added and centrifuged at 380 × *g* for 5 min, repeatedly. The supernatant containing crude mitochondria was centrifuged at 4,500 × *g* for 5 min and resulting pellet was resuspended and frozen at -80°C until use. The pellets were sonicated and diluted with Mitochondrial extraction buffer (Abcam, MA, USA) for protein determination by BCA assay (Pierce, IL, USA). The mitochondrial respiratory chain complex (MRCC) enzymes from the isolated mitochondria were determined with MitoTox complete OXPHOS activity assay kit (Abcam, MA, USA): rotenone-sensitive NADH-ubiquinone oxidoreductase (MRCC I), succinate dehydrogenase (MRCC II), antimycin A-sensitive decylubiquinol cytochrome c oxidoreductase (MRCC III), cytochrome c oxidase (MRCC IV), and ATP synthase (MRCC V). Briefly, frozen samples of isolated mitochondria were thawed on ice and solubilized with 2% CHAPS and aliquots were used for enzymatic activity assay following the protocol of the manufacturer. The reactions used for the determination of MRCC were monitored using fluorescence plate reader (Synergy H1, BioTek, VT, USA) normalization with total protein concentration (BCA assay, Pierce, IL, USA) as reported in nmol substrate/min/mg protein.

#### Measurement of ATP generation

Generated ATP levels were quantified using colorimetric method based ATP assay kit (Abcam, MA, USA). Briefly, synchronized worms were collected and washed with M9 buffer. The worm pellets were treated freeze and thaw cycles and boiled for 15 min to release ATP and inactive ATPase. Pellets were centrifuged at 12,000 × *g* for 10 min and supernatant were used for the assay following instructions normalization with total protein concentration (BCA assay, Pierce, IL, USA).

#### ATFS-1 and SKN-1 localization assay

The *skn-1*(*zu169*)*IV*;*geIs7*(LG326), *skn-1*(*zu135*)*IV*;*gels7*(LG333), and *wgIs675*[*atfs-1::TY1::EGFP::3xFLAG* + *unc-119*(+)](OP675) strain was used and their fluorescence was visualized using a confocal laser scanning microscope with inverted stand (LSM8, Carl Zeiss, Jena, Germany) (with excitation at 488 nm and emission at 535 nm). For each condition, the nucleus/intermediate/cytoplasm fluorescence or fluorescence intensity of 30 - 40 worms were detected densitometrically with ZEN Lite software (Carl Zeiss, Jena, Germany) with triplicate attempts.

#### Bioinformatics in skn-1, atfs-1 RNAi public data

Data pertaining to *skn-1* RNAi and *atfs-1* RNAi experiments were retrieved from the NCBI GEO database, specifically GSE63075 and GSE179517, respectively. Differential expression gene analysis was conducted using DESeq2 version 1.38.3 as previously described^61^.

#### Human study participants

This was a cross-sectional study performed on a cohort of ambulatory, community-dwelling older adults. The study population comprised Koreans who had undergone a comprehensive geriatric assessment at the Division of Endocrinology and Metabolism, Department of Internal Medicine, Chungnam National University Hospital (CNUH), between April 2021 and April 2023. These participants visited the clinic for the management of chronic diseases, such as hypertension, dyslipidemia, osteoporosis, and diabetes, or for the evaluation of nonspecific symptoms, such as fatigue and loss of appetite, which are frequently observed in older adults. They were not from nursing homes or inpatient facilities and were consecutively enrolled. The exclusion criteria were as follows: a history of ongoing cancer or a cancer-free period <5 years; psychiatric disease; malabsorption disorder (including diabetic gastropathy or a history of bowel resection); pregnancy; acute bacterial or viral infection; or requirement of a wheelchair for mobility.

#### Ethical considerations

All participants received information about the goals and procedures of the study, and all agreed to participate by signing a consent form. The study was approved by the Institutional Review Boards of Chungnam National University Hospital (2019-06-063-016), and written informed consent was obtained from all enrolled participants.

#### Sarcopenia assessment

Information on demographic characteristics and medical and surgical histories was collected through detailed interviews and reviews of medical records by experienced nurses. Body composition including muscle mass (whole body lean body mass minus bone mineral content) was evaluated using a bioelectrical impedance analyzer (InBody 270 Body Composition Analyzer; InBody USA, Cerritos, CA, USA) with measuring frequencies of 1, 5, 50, 250, 500, and 1000 kHz^69^. Appendicular skeletal muscle mass (ASM) was defined as the sum of the muscle mass of all four limbs. The skeletal muscle index (SMI) was defined as appendicular skeletal muscle mass divided by height squared (ASM/m^2^)^70^. Handgrip strength of the dominant side was measured using a Smedley type dynamometer (Takei T.K.K.5401 GRIP-D handgrip dynamometer; Takei Scientific Instruments Co., Ltd, Tokyo, Japan). Participants were instructed to adopt a comfortable sitting position, bend their elbows to 90° (90° flexion), and squeeze the dynamometer as hard as possible. The maximum value was adjusted after all tests were conducted twice at 1-min intervals or longer. Physical performance was evaluated using the short physical performance battery (SPPB), which includes standardized performance tests: gait speed in a 4-m walk test; a five times sit-to stand test (5TSTS) of coordination and strength; and a tandem test for static balance. The tandem test was performed in three different positions: a side-by-side position, a semi-tandem position, and a full tandem position. The participants were asked to maintain each position for >10 s, and the amount of time (s) that they successfully remained in the given position was recorded. The higher the SPPB score (which ranged from 0 to 12 points), the better the lower extremity function. A diagnosis of sarcopenia was based on the 2019 Consensus Guidelines from the Asian Working Group for Sarcopenia^71,72^. Patients with low appendicular muscle mass (SMI measured by bioelectrical impedance analyzer: <7.0 kg/m^2^ for men and <5.7 kg/m^2^ for women) and low muscle strength (handgrip strength <28 kg for men and <18 kg for women) with or without low physical performance (gait speed <1.0 m/s; 5TSTS ≥12 s; or SPPB score ≤9 points) were classified as having sarcopenia.

#### Serum Metabolites Extraction

Metabolites were extracted using a modified protocol from previous publications.^73,74^ Briefly, 150 μL of human serum was transferred into an Eppendorf tube containing 450 μL of MS-grade methanol supplemented with 100uM DL-Norvaline as an internal standard. 200 μL of chloroform were subsequently added into the samples, and thoroughly vortexed for 30 s. The samples were then centrifuged for 20 mins at 15,000 g at 4 L following addition of 200 μL of MS-grade water. 700 μL of the aqueous phase was carefully transferred into a new microcentrifuge tube, followed by 6 h of drying in vacuum centrifuge at 4 L. The completely dried pellet was dissolved in 20 μL of 50 % methanol, and transferred into a glass insert for the subsequent LC-MS analysis.

#### Measurement of D-3-Phenyllactic acid Using LC-QTOF

The metabolite in human serum was analyzed on Agilent 1290 infinity II LC system coupled to Agilent Poroshell 120 HILIC-Z (3 x 150 mm, 2.7 μm) and Agilent 6530 quadrupole time of flight mass spectrometer using Dual Agilent Jet Stream Electrospary Ionization. Injection volume of sample was 5 μL. LC separation was achieved on Agilent Poroshell 120 HILIC-Z column(3 x 150 mm, 2.7 μm) using mobile phase A(10 mM ammonium acetate in water, pH 9.0) and mobile phase B(10 mM ammonium acetate water/acetonitrile 15:85 (v:v) pH 9.0). Flow rate was 0.300 mL/min. The LC gradient was set at ****0 min 95% B, 2 min 95% B, 10 min 70% B, 12 min 60% B, 14 min 40% min, 17 min 40% B, 19 min 95% B. Electrospary ionization parameters included gas temperature at 300 L, drying gas flow of 10 L/min, nebulizer pressure of 40 psi, sheath gas temperature at 350 L, sheath gas flow of 12 L/min, capillary voltage of 3000 V, fragmentor of 120 V, skimmer of 65 V, Oct 1 RF vpp of 750 V. Compound were detected using electrospray ionization source operating under extended dynamic range(EDR 1700 m/z) in negative ionization mode. Peak area integration was performed using MassHunter Q-TOF Quantitative Analysis (Agilent). Peak identification of D-Phenyllactic acid in each sample was carried out based on the comparison of the retention time and accurate mass-tocharge ratio from D-Phenyllactic acid (Sigma-Aldrich, MO, USA) analyzed under the identical conditions.

#### PLA analysis in human subjects

To compare plasma PLA levels between age-matched elderly individuals and those with sarcopenia, we used the result of LC-QTOF and indicators associated with sarcopenia, such as grip strength, gait speed, 5-time sit-to-stand, and SPPB score. We generated boxplots with Rstudio (RStudio Desktop 1.4.1717; R 4.1.1) installed R packages dplyr, stringr, ggpubr, ggplot2, pheatmap, igraph, ggraph, corrr, corrplot, tidyverse, and reshape2 and carried out statistical analysis comparing two groups, non-sarcopenic and sarcopenic as previously described^75^. We employed one-tailed t-tests for the analysis.

#### Statistical analysis

All experiments were repeated at least three times with identical or similar results. Data represents biological replicates. Adequate statistical analysis was used for every assays. Data meet the hypothesis of the statistical tests described each experiment. Data are expressed as the mean ± s.d. in all figures unless stated otherwise. R software (ver. 4.1.1) was used for statistical analyses. The *p*-values \**p* < 0.05, \*\**p* < 0.01, \*\*\**p* < 0.001, and ^#^*p* < 0.01 were considered statistically significant.

#### Data availability

There is no restriction on data availability. The RNA sequence data used in this study have been deposited in NCBI’s SRA and are accessible through submission number PRJNA827000 and PRJNA894441. Source data for all the individual *P* values are provided with this paper.

## References

1. Weyh, C., Krüger, K., and Strasser, B. (2020). Physical Activity and Diet Shape the Immune System during Aging. Nutrients 12. 10.3390/nu12030622.

2. Gimeno-Mallench, L., Sanchez-Morate, E., Parejo-Pedrajas, S., Mas-Bargues, C., Inglés, M., Sanz-Ros, J., Román-Domínguez, A., Olaso, G., Stromsnes, K., and Gambini, J. (2020). The Relationship between Diet and Frailty in Aging. Endocrine, metabolic & immune disorders drug targets 20, 1373–1382. 10.2174/1871530320666200513083212.

3. Fontana, L., and Partridge, L. (2015). Promoting health and longevity through diet: from model organisms to humans. Cell 161, 106–118. 10.1016/j.cell.2015.02.020.

4. Li, W., Gao, L., Huang, W., Ma, Y., Muhammad, I., Hanif, A., Ding, Z., and Guo, X. (2022). Antioxidant properties of lactic acid bacteria isolated from traditional fermented yak milk and their probiotic effects on the oxidative senescence of Caenorhabditis elegans. Food & function 13, 3690–3703. 10.1039/d1fo03538j.

5. Liu, G., Tan, F.H., Lau, S.A., Jaafar, M.H., Chung, F.Y., Azzam, G., Liong, M.T., and Li, Y. (2022). Lactic acid bacteria feeding reversed the malformed eye structures and ameliorated gut microbiota profiles of Drosophila melanogaster Alzheimer’s disease model. Journal of applied microbiology 132, 3155–3167. 10.1111/jam.14773.

6. Jin, X., He, Y., Zhou, Y., Chen, X., Lee, Y.K., Zhao, J., Zhang, H., Chen, W., and Wang, G. (2021). Lactic acid bacteria that activate immune gene expression in Caenorhabditis elegans can antagonise Campylobacter jejuni infection in nematodes, chickens and mice. BMC microbiology 21, 169. 10.1186/s12866-021-02226-x.

7. Schretter, C.E., Vielmetter, J., Bartos, I., Marka, Z., Marka, S., Argade, S., and Mazmanian, S.K. (2018). A gut microbial factor modulates locomotor behaviour in Drosophila. Nature 563, 402–406. 10.1038/s41586-018-0634-9.

8. De Filippis, F., Pasolli, E., and Ercolini, D. (2020). The food-gut axis: lactic acid bacteria and their link to food, the gut microbiome and human health. FEMS microbiology reviews 44, 454–489. 10.1093/femsre/fuaa015.

9. Markowiak-Kopeć, P., and Śliżewska, K. (2020). The Effect of Probiotics on the Production of Short-Chain Fatty Acids by Human Intestinal Microbiome. Nutrients 12. 10.3390/nu12041107.

10. Dalile, B., Van Oudenhove, L., Vervliet, B., and Verbeke, K. (2019). The role of short-chain fatty acids in microbiota-gut-brain communication. Nature reviews. Gastroenterology & hepatology 16, 461–478. 10.1038/s41575-019-0157-3.

11. Morrison, D.J., and Preston, T. (2016). Formation of short chain fatty acids by the gut microbiota and their impact on human metabolism. Gut microbes 7, 189–200. 10.1080/19490976.2015.1134082.

12. Koh, A., De Vadder, F., Kovatcheva-Datchary, P., and Bäckhed, F. (2016). From Dietary Fiber to Host Physiology: Short-Chain Fatty Acids as Key Bacterial Metabolites. Cell 165, 1332–1345. 10.1016/j.cell.2016.05.041.

13. Huang, W., Man, Y., Gao, C., Zhou, L., Gu, J., Xu, H., Wan, Q., Long, Y., Chai, L., Xu, Y., and Xu, Y. (2020). Short-Chain Fatty Acids Ameliorate Diabetic Nephropathy via GPR43-Mediated Inhibition of Oxidative Stress and NF-κB Signaling. Oxidative medicine and cellular longevity 2020, 4074832. 10.1155/2020/4074832.

14. Chae, M., Kim, B.J., Na, J., Kim, S.Y., Lee, J.O., Kim, Y.J., Lee, E., Cho, D., Roh, J., and Kim, W. (2021). Antimicrobial activity of Lactiplantibacillus plantarum APsulloc 331261 and APsulloc 331266 against pathogenic skin microbiota. Frontiers in bioscience (Elite edition) 13, 237–248. 10.52586/e881.

15. Arellano, K., Vazquez, J., Park, H., Lim, J., Ji, Y., Kang, H.J., Cho, D., Jeong, H.W., and Holzapfel, W.H. (2020). Safety Evaluation and Whole-Genome Annotation of Lactobacillus plantarum Strains from Different Sources with Special Focus on Isolates from Green Tea. Probiotics and antimicrobial proteins 12, 1057–1070. 10.1007/s12602-019-09620-y.

16. Park, H., Cho, D., Huang, E., Seo, J.Y., Kim, W.G., Todorov, S.D., Ji, Y., and Holzapfel, W.H. (2020). Amelioration of Alcohol Induced Gastric Ulcers Through the Administration of Lactobacillus plantarum APSulloc 331261 Isolated From Green Tea. Frontiers in microbiology 11, 420. 10.3389/fmicb.2020.00420.

17. Wei, M., Brandhorst, S., Shelehchi, M., Mirzaei, H., Cheng, C.W., Budniak, J., Groshen, S., Mack, W.J., Guen, E., Di Biase, S., et al. (2017). Fasting-mimicking diet and markers/risk factors for aging, diabetes, cancer, and cardiovascular disease. Science translational medicine 9. 10.1126/scitranslmed.aai8700.

18. Maki, Y., Soejima, H., Kitamura, T., Sugiyama, T., Sato, T., Watahiki, M.K., and Yamaguchi, J. (2021). 3-Phenyllactic acid, a root-promoting substance isolated from Bokashi fertilizer, exhibits synergistic effects with tryptophan. Plant biotechnology (Tokyo, Japan) 38, 9–16. 10.5511/plantbiotechnology.20.0727a.

19. Long, D.M., Frame, A.K., Reardon, P.N., Cumming, R.C., Hendrix, D.A., Kretzschmar, D., and Giebultowicz, J.M. (2020). Lactate dehydrogenase expression modulates longevity and neurodegeneration in Drosophila melanogaster. Aging 12, 10041–10058. 10.18632/aging.103373.

20. Tauffenberger, A., Fiumelli, H., Almustafa, S., and Magistretti, P.J. (2019). Lactate and pyruvate promote oxidative stress resistance through hormetic ROS signaling. Cell death & disease 10, 653. 10.1038/s41419-019-1877-6.

21. Li, P., Lu, Y., Zhao, M., Chen, L., Zhang, C., Cheng, Q., and Chen, C. (2021). Effects of Phenyllactic Acid, Lactic Acid Bacteria, and Their Mixture on Fermentation Characteristics and Microbial Community Composition of Timothy Silage. Frontiers in microbiology 12, 743433. 10.3389/fmicb.2021.743433.

22. Mu, W., Yu, S., Zhu, L., Zhang, T., and Jiang, B. (2012). Recent research on 3-phenyllactic acid, a broad-spectrum antimicrobial compound. Applied microbiology and biotechnology 95, 1155–1163. 10.1007/s00253-012-4269-8.

23. Saul, N., Moller, S., Cirulli, F., Berry, A., Luyten, W., and Fuellen, G. (2021). Health and longevity studies in C. elegans: the “healthy worm database” reveals strengths, weaknesses and gaps of test compound-based studies. Biogerontology 22, 215–236. 10.1007/s10522-021-09913-2.

24. Rabinowitz, J.D., and Enerbäck, S. (2020). Lactate: the ugly duckling of energy metabolism. Nature metabolism 2, 566–571. 10.1038/s42255-020-0243-4.

25. Brooks, G.A. (2018). The Science and Translation of Lactate Shuttle Theory. Cell metabolism 27, 757–785. 10.1016/j.cmet.2018.03.008.

26. Goodpaster, B.H., and Sparks, L.M. (2017). Metabolic Flexibility in Health and Disease. Cell metabolism 25, 1027–1036. 10.1016/j.cmet.2017.04.015.

27. Hui, S., Ghergurovich, J.M., Morscher, R.J., Jang, C., Teng, X., Lu, W., Esparza, L.A., Reya, T., Le, Z., Yanxiang Guo, J., et al. (2017). Glucose feeds the TCA cycle via circulating lactate. Nature 551, 115–118. 10.1038/nature24057.

28. Kitaoka, Y., Ogborn, D.I., Mocellin, N.J., Schlattner, U., and Tarnopolsky, M.A. (2013). Monocarboxylate transporters and mitochondrial creatine kinase protein content in McArdle disease. Molecular genetics and metabolism 108, 259–262. 10.1016/j.ymgme.2013.01.005.

29. Possemiers, H., Vandermosten, L., and Van den Steen, P.E. (2021). Etiology of lactic acidosis in malaria. PLoS pathogens 17, e1009122. 10.1371/journal.ppat.1009122.

30. Kraut, J.A., and Madias, N.E. (2014). Lactic acidosis. The New England journal of medicine 371, 2309–2319. 10.1056/NEJMra1309483.

31. Vercellino, I., and Sazanov, L.A. (2022). The assembly, regulation and function of the mitochondrial respiratory chain. Nature reviews. Molecular cell biology 23, 141–161. 10.1038/s41580-021-00415-0.

32. Takahashi, K., Tamura, Y., Kitaoka, Y., Matsunaga, Y., and Hatta, H. (2022). Effects of Lactate Administration on Mitochondrial Respiratory Function in Mouse Skeletal Muscle. Frontiers in physiology 13, 920034. 10.3389/fphys.2022.920034.

33. Maglioni, S., Mello, D.F., Schiavi, A., Meyer, J.N., and Ventura, N. (2019). Mitochondrial bioenergetic changes during development as an indicator of C. elegans health-span. Aging 11, 6535–6554. 10.18632/aging.102208.

34. Macedo, F., Romanatto, T., Gomes de Assis, C., Buis, A., Kowaltowski, A.J., Aguilaniu, H., and Marques da Cunha, F. (2020). Lifespan-extending interventions enhance lipid-supported mitochondrial respiration in Caenorhabditis elegans. FASEB journal : official publication of the Federation of American Societies for Experimental Biology 34, 9972–9981. 10.1096/fj.201901880R.

35. Wu, Z., Senchuk, M.M., Dues, D.J., Johnson, B.K., Cooper, J.F., Lew, L., Machiela, E., Schaar, C.E., DeJonge, H., Blackwell, T.K., and Van Raamsdonk, J.M. (2018). Mitochondrial unfolded protein response transcription factor ATFS-1 promotes longevity in a long-lived mitochondrial mutant through activation of stress response pathways. BMC biology 16, 147. 10.1186/s12915-018-0615-3.

36. Nargund, A.M., Pellegrino, M.W., Fiorese, C.J., Baker, B.M., and Haynes, C.M. (2012). Mitochondrial import efficiency of ATFS-1 regulates mitochondrial UPR activation. Science 337, 587–590. 10.1126/science.1223560.

37. Organization, W.H. (2021). Decade of healthy ageing: baseline report. https://iris.who.int/bitstream/handle/10665/341488/9789240023307-eng.pdf.

38. Nasrollahzadeh, A., Mokhtari, S., Khomeiri, M., and Saris, P.E.J. (2022). Antifungal Preservation of Food by Lactic Acid Bacteria. Foods (Basel, Switzerland) 11. 10.3390/foods11030395.

39. Daniel, C., Roussel, Y., Kleerebezem, M., and Pot, B. (2011). Recombinant lactic acid bacteria as mucosal biotherapeutic agents. Trends in biotechnology 29, 499–508. 10.1016/j.tibtech.2011.05.002.

40. Kim, H., Kim, M., Myoung, K., Kim, W., Ko, J., Kim, K.P., and Cho, E.G. (2020). Comparative Lipidomic Analysis of Extracellular Vesicles Derived from Lactobacillus plantarum APsulloc 331261 Living in Green Tea Leaves Using Liquid Chromatography-Mass Spectrometry. International journal of molecular sciences 21. 10.3390/ijms21218076.

41. Yanase, S., Yasuda, K., and Ishii, N. (2019). Monitoring Age-Related Changes in the Lactate/Pyruvate Ratio Using a Colorimetric Assay in a C. elegans Model of Increased Life Span. Methods in molecular biology (Clifton, N.J.) 1916, 123–132. 10.1007/978-1-4939-8994-2_12.

42. Grad, L.I., Sayles, L.C., and Lemire, B.D. (2005). Introduction of an additional pathway for lactate oxidation in the treatment of lactic acidosis and mitochondrial dysfunction in Caenorhabditis elegans. Proceedings of the National Academy of Sciences of the United States of America 102, 18367–18372. 10.1073/pnas.0506939102.

43. Cairns, S.P. (2006). Lactic acid and exercise performance : culprit or friend? Sports medicine (Auckland, N.Z.) 36, 279–291. 10.2165/00007256-200636040-00001.

44. Brooks, G.A. (2020). Lactate as a fulcrum of metabolism. Redox biology 35, 101454. 10.1016/j.redox.2020.101454.

45. Faubert, B., Li, K.Y., Cai, L., Hensley, C.T., Kim, J., Zacharias, L.G., Yang, C., Do, Q.N., Doucette, S., Burguete, D., et al. (2017). Lactate Metabolism in Human Lung Tumors. Cell 171, 358–371.e359. 10.1016/j.cell.2017.09.019.

46. Huang, Y., Zhang, J., Dong, R., Ji, X., Jiang, Y., Cen, J., Bai, Z., Hong, K., Li, H., Chen, J., et al. (2021). Lactate as a metabolite from probiotic Lactobacilli mitigates ethanol-induced gastric mucosal injury: an in vivo study. BMC complementary medicine and therapies 21, 26. 10.1186/s12906-020-03198-7.

47. Saez-Lara, M.J., Gomez-Llorente, C., Plaza-Diaz, J., and Gil, A. (2015). The role of probiotic lactic acid bacteria and bifidobacteria in the prevention and treatment of inflammatory bowel disease and other related diseases: a systematic review of randomized human clinical trials. BioMed research international 2015, 505878. 10.1155/2015/505878.

48. Lee, Y.S., Kim, T.Y., Kim, Y., Lee, S.H., Kim, S., Kang, S.W., Yang, J.Y., Baek, I.J., Sung, Y.H., Park, Y.Y., et al. (2018). Microbiota-Derived Lactate Accelerates Intestinal Stem-Cell-Mediated Epithelial Development. Cell host & microbe 24, 833–846.e836. 10.1016/j.chom.2018.11.002.

49. Jang, C., Hui, S., Zeng, X., Cowan, A.J., Wang, L., Chen, L., Morscher, R.J., Reyes, J., Frezza, C., Hwang, H.Y., et al. (2019). Metabolite Exchange between Mammalian Organs Quantified in Pigs. Cell metabolism 30, 594–606.e593. 10.1016/j.cmet.2019.06.002.

50. van Gemert, L.A., de Galan, B.E., Wevers, R.A., Ter Heine, R., and Willemsen, M.A. (2022). Lactate infusion as therapeutical intervention: a scoping review. European journal of pediatrics. 10.1007/s00431-022-04446-3.

51. Salekeen, R., Siam, M.H.B., Sharif, D.I., Lustgarten, M.S., Billah, M.M., and Islam, K.M.D. (2021). In silico insights into potential gut microbial modulation of NAD+ metabolism and longevity. Journal of biochemical and molecular toxicology 35, e22925. 10.1002/jbt.22925.

52. Birch, J., and Gil, J. (2020). Senescence and the SASP: many therapeutic avenues. Genes & development 34, 1565–1576. 10.1101/gad.343129.120.

53. Vatner, S.F., Zhang, J., Oydanich, M., Berkman, T., Naftalovich, R., and Vatner, D.E. (2020). Healthful aging mediated by inhibition of oxidative stress. Ageing research reviews 64, 101194. 10.1016/j.arr.2020.101194.

54. Campisi, J., Kapahi, P., Lithgow, G.J., Melov, S., Newman, J.C., and Verdin, E. (2019). From discoveries in ageing research to therapeutics for healthy ageing. Nature 571, 183–192. 10.1038/s41586-019-1365-2.

55. Sims, C.A., Labiner, H.E., Shah, S.S., and Baur, J.A. (2021). Longevity pathways in stress resistance: targeting NAD and sirtuins to treat the pathophysiology of hemorrhagic shock. GeroScience 43, 1217–1228. 10.1007/s11357-020-00311-z.

56. Salminen, A., Kaarniranta, K., and Kauppinen, A. (2017). Integrated stress response stimulates FGF21 expression: Systemic enhancer of longevity. Cellular signalling 40, 10–21. 10.1016/j.cellsig.2017.08.009.

57. Vintila, A.R., Slade, L., Cooke, M., Willis, C.R.G., Torregrossa, R., Rahman, M., Anupom, T., Vanapalli, S.A., Gaffney, C.J., Gharahdaghi, N., et al. (2023). Mitochondrial sulfide promotes life span and health span through distinct mechanisms in developing versus adult treated Caenorhabditis elegans. Proc Natl Acad Sci U S A 120, e2216141120. 10.1073/pnas.2216141120.

58. Tullet, J.M., Hertweck, M., An, J.H., Baker, J., Hwang, J.Y., Liu, S., Oliveira, R.P., Baumeister, R., and Blackwell, T.K. (2008). Direct inhibition of the longevity-promoting factor SKN-1 by insulin-like signaling in C. elegans. Cell 132, 1025–1038. 10.1016/j.cell.2008.01.030.

59. Blackwell, T.K., Steinbaugh, M.J., Hourihan, J.M., Ewald, C.Y., and Isik, M. (2015). SKN-1/Nrf, stress responses, and aging in Caenorhabditis elegans. Free Radic Biol Med 88, 290–301. 10.1016/j.freeradbiomed.2015.06.008.

60. Ravitch, M.M. (1969). Spontaneous pneumothorax. Med Times 97, 274–276.

61. Kim, J., Jo, Y., Cho, D., and Ryu, D. (2022). L-threonine promotes healthspan by expediting ferritin-dependent ferroptosis inhibition in C. elegans. Nat Commun 13, 6554. 10.1038/s41467-022-34265-x.

62. Hibshman, J.D., Webster, A.K., and Baugh, L.R. (2021). Liquid-culture protocols for synchronous starvation, growth, dauer formation, and dietary restriction of Caenorhabditis elegans. STAR protocols 2, 100276. 10.1016/j.xpro.2020.100276.

63. Dobin, A., Davis, C.A., Schlesinger, F., Drenkow, J., Zaleski, C., Jha, S., Batut, P., Chaisson, M., and Gingeras, T.R. (2013). STAR: ultrafast universal RNA-seq aligner. Bioinformatics (Oxford, England) 29, 15–21. 10.1093/bioinformatics/bts635.

64. Bolger, A.M., Lohse, M., and Usadel, B. (2014). Trimmomatic: a flexible trimmer for Illumina sequence data. Bioinformatics (Oxford, England) 30, 2114–2120. 10.1093/bioinformatics/btu170.

65. Pertea, M., Pertea, G.M., Antonescu, C.M., Chang, T.C., Mendell, J.T., and Salzberg, S.L. (2015). StringTie enables improved reconstruction of a transcriptome from RNA-seq reads. Nature biotechnology 33, 290–295. 10.1038/nbt.3122.

66. Pertea, M., Kim, D., Pertea, G.M., Leek, J.T., and Salzberg, S.L. (2016). Transcript-level expression analysis of RNA-seq experiments with HISAT, StringTie and Ballgown. Nature protocols 11, 1650–1667. 10.1038/nprot.2016.095.

67. Kim, W., Underwood, R.S., Greenwald, I., and Shaye, D.D. (2018). OrthoList 2: A New Comparative Genomic Analysis of Human and Caenorhabditis elegans Genes. Genetics 210, 445–461. 10.1534/genetics.118.301307.

68. Hoogewijs, D., Houthoofd, K., Matthijssens, F., Vandesompele, J., and Vanfleteren, J.R. (2008). Selection and validation of a set of reliable reference genes for quantitative sod gene expression analysis in C. elegans. BMC molecular biology 9, 9. 10.1186/1471-2199-9-9.

69. Oh, J.H., Song, S., Rhee, H., Lee, S.H., Kim, D.Y., Choe, J.C., Ahn, J., Park, J.S., Shin, M.J., Jeon, Y.K., et al. (2019). Normal Reference Plots for the Bioelectrical Impedance Vector in Healthy Korean Adults. J Korean Med Sci 34, e198. 10.3346/jkms.2019.34.e198.

70. Nga, H.T., Jang, I.Y., Kim, D.A., Park, S.J., Lee, J.Y., Lee, S., Kim, J.H., Lee, E., Park, J.H., Lee, Y.H., et al. (2021). Serum GDF15 Level Is Independent of Sarcopenia in Older Asian Adults. Gerontology 67, 525–531. 10.1159/000513600.

71. Chen, L.K., Woo, J., Assantachai, P., Auyeung, T.W., Chou, M.Y., Iijima, K., Jang, H.C., Kang, L., Kim, M., Kim, S., et al. (2020). Asian Working Group for Sarcopenia: 2019 Consensus Update on Sarcopenia Diagnosis and Treatment. J Am Med Dir Assoc 21, 300–307 e302. 10.1016/j.jamda.2019.12.012.

72. Dao, T., Green, A.E., Kim, Y.A., Bae, S.J., Ha, K.T., Gariani, K., Lee, M.R., Menzies, K.J., and Ryu, D. (2020). Sarcopenia and Muscle Aging: A Brief Overview. Endocrinol Metab (Seoul) 35, 716–732. 10.3803/EnM.2020.405.

73. Dodd, D., Spitzer, M.H., Van Treuren, W., Merrill, B.D., Hryckowian, A.J., Higginbottom, S.K., Le, A., Cowan, T.M., Nolan, G.P., Fischbach, M.A., and Sonnenburg, J.L. (2017). A gut bacterial pathway metabolizes aromatic amino acids into nine circulating metabolites. Nature 551, 648–652. 10.1038/nature24661.

74. Molenaars, M., Schomakers, B.V., Elfrink, H.L., Gao, A.W., Vervaart, M.A.T., Pras-Raves, M.L., Luyf, A.C., Smith, R.L., Sterken, M.G., Kammenga, J.E., et al. (2021). Metabolomics and lipidomics in Caenorhabditis elegans using a single-sample preparation. Dis Model Mech 14. 10.1242/dmm.047746.

75. Kim, J., Lee, H., Jin, E.J., Jo, Y., Kang, B.E., Ryu, D., and Kim, G. (2022). A Microfluidic Device to Fabricate One-Step Cell Bead-Laden Hydrogel Struts for Tissue Engineering. Small 18, e2106487. 10.1002/smll.202106487.

